# Synanthropic flies harbor a distinct, pathogen-enriched microbial compartment in livestock landscapes

**DOI:** 10.64898/2026.08.05.743036

**Authors:** Alicia Montemayor, Phillip E. Kaufman, T. Matthew Taylor, Giridhar Athrey

**Author notes:** **Corresponding Author** Giri Athrey.

## Abstract

Zoonotic and other livestock-associated pathogens move readily within animal production systems, yet their dispersal into surrounding environments is poorly characterized. We evaluated whether synanthropic filth flies act as spatially integrating sentinels of livestock-associated pathobiomes. From 30 roadside sites in six Texas counties stratified by livestock production intensity, we collected 3,373 Calliphoridae spp. and 12 *Musca domestica*, together with co-located soil, water, plant, and fecal samples, and characterized 195 metagenomes (122 insect, 73 environmental). Using diversity, differential-abundance, machine-learning, indicator-species, and network analyses, we found three robust patterns. First, fly metagenomes form a microbial compartment compositionally distinct from co-located environmental matrices (PERMANOVA R² = 0.094, F = 19.81, P = 0.001) and carry markedly higher relative abundances of pathogen-associated taxa, so insect sampling is complementary to, not redundant with, environmental sampling. Second, the dominant landscape signal is categorical rather than graded: cattle presence, not a Low–Medium–High density gradient, most strongly correlated with community structure, with the ruminant-associated anaerobe *Peptostreptococcus russellii* as the most reproducible biomarker. Third, intensification produced taxon-level enrichments (notably the feedlot pathogen *Trueperella pyogenes*) and progressive co-occurrence-network fragmentation, without a monotonic dose-response. A curated 12-pathogen panel failed to classify landscape context, whereas whole-community analysis succeeded. Filth flies thus provide a pragmatic, spatially integrated readout that distinguishes low-impact from livestock-impacted landscapes, offering a defensible tool for One Health pathogen surveillance.

**IMPORTANCE:** Zoonotic and other livestock-associated pathogens emerging from animal production are among the most pressing threats to global health but tracking how they spill into the wider environment is difficult and expensive. We show that common filth flies, abundant, easy to trap, and free ranging across farms, fields, and human spaces, act as living samplers that concentrate and report the microbial signature of nearby livestock. Sampling flies captured information that soil and water sampling missed, and a single binary signal, the presence or absence of cattle, was associated with the microbial community more strongly than any measure of animal density. Narrow tests targeting a short curated list of pathogens were not successful in differentiating sites, while analyzing the whole microbial community provided more discriminating power. These findings support a practical, low-cost strategy in which insects serve as biological sentinels for the detection of livestock-associated pathogens across agricultural landscapes.

## INTRODUCTION

Antimicrobial resistance (AMR) is a defining One Health challenge, with resistant infections projected to impose severe economic and public-health costs over the coming decades (1, 2). Livestock production is a major node in the emergence and dissemination of resistant and zoonotic organisms, yet pathogens are typically well documented only within farm boundaries; their spatial dynamics and dispersal into surrounding soils, waters, and human-inhabited spaces remain poorly resolved (3, 4). Environmental reservoirs (i.e., water, air, soil) are increasingly recognized as conduits for resistance, but point-source sampling of these matrices is logistically demanding and captures only a static, spatially restricted snapshot (5, 6).

The selection pressure created by antimicrobial use in food-animal production is transmitted to surrounding ecosystems through manure application, runoff, aerosols, and contaminated water, seeding environmental reservoirs that can return resistance to humans and animals (5, 6, 8). Resistance genes and resistant organisms have been traced from above-ground livestock sources into groundwater and along the farm-to-urban interface, demonstrating that the resistome is not confined to the operation that generates it. Yet capturing this dispersal empirically is difficult; soil, water, and fecal samples are spatially fixed, vary enormously over short distances, and require dense grids of collection points to approximate landscape-scale exposure. A sampling unit that physically integrates the microbial environment across space would substantially reduce this burden.

Synanthropic filth flies (Calliphoridae and Muscidae) are promising complements to environmental sampling. Their developmental and feeding ecology brings them into repeated contact with manure, carrion, wounds, and secretions, and their mobility integrates exposures across a foraging range of roughly one to two kilometers (7). House flies and blow flies are known to harbor and disseminate antimicrobial-resistant and enteropathogenic bacteria in agricultural, hospital, and urban settings, including prior work on Texas farms (8–13, 35), and fly-associated microbiomes form characteristic assemblages that segregate by host species and habitat (14). Flies have additionally been implicated as carriers of zoonotic pathogens on dairy farms (13) and even as vehicles for highly pathogenic avian influenza virus near affected premises (37), reinforcing their relevance to One Health surveillance. Most prior studies, however, rely on culture or targeted assays focused on a small number of indicator organisms. Few have used whole-community shotgun metagenomics to ask what fraction of the livestock pathobiome flies actually capture relative to the co-located environment they sample, or whether the fly signal is quantitatively graded or is categorically threshold-like with respect to livestock pressure.

We frame filth flies as bioaccumulating sentinels, by analogy to aquatic bivalves used to monitor bioavailable contaminants: organisms that integrate exposure over space and time and concentrate a biologically filtered signal. We tested this concept through whole-microbiome metagenomics of pooled flies collected along a stratified livestock intensity gradient in Texas, alongside co-located environmental samples. Our objectives were to (i) determine whether insect-derived microbiomes form a compositionally distinct compartment relative to the environment; (ii) identify the landscape drivers, livestock intensity, cattle presence, and geography, that structure that compartment; (iii) detect pathogens, indicator taxa, and community-level structural changes that respond to livestock pressure; and (iv) evaluate whether a narrow targeted pathogen panel can substitute for whole-microbiome analysis in classifying landscape context. Four focal hypotheses structured the analysis, distinguishing smooth dose-response expectations from categorical ecological contrasts. Because livestock intensity and cattle presence were assigned at the county scale, the landscape comparisons are interpreted as observational associations across six counties.

## RESULTS

### Fly metagenomes form a distinct, pathogen-enriched compartment

We collected 3,373 Calliphoridae spp. and 12 *Musca domestica* across 30 sites in six counties; Calliphoridae accounted for 99.6% of specimens, with the highest yield in Coryell County and the lowest in Gonzales County (Table 1). After prevalence filtering, 195 metagenomes (122 insect, 73 environmental) and 254 taxa were retained for analysis.

**TABLE 1.** Total abundance and taxonomic distribution of flies collected across the six sampling counties.

|  | Grimes<br>(Low) | Washington<br>(Low) | Coryell<br>(Med) | Bastrop<br>(Med) | Gonzales<br>(High) | Shelby<br>(High) | Total |
| --- | --- | --- | --- | --- | --- | --- | --- |
| *Calliphoridae*<br>*spp.* | 787 | 366 | 1123 | 362 | 315 | 420 | 3,373 |
| *Musca*<br>*domestica* | 0 | 3 | 0 | 2 | 4 | 3 | 12 |

The strongest single effect in the dataset was the compositional separation of insect-derived microbiomes from co-located environmental samples (PERMANOVA R² = 0.094, F = 19.81, P = 0.001; Fig. 1A). Environmental samples were also significantly more diverse (Wilcoxon P < 0.001; Fig. S1), consistent with the fly gut acting as a selective filter rather than a passive mirror of its substrate. Critically, insects carried dramatically higher relative abundances of pathogen-associated taxa than the environment (Wilcoxon P < 2.2 × 10⁻¹⁶, lower bound; Fig. 1B). Most environmental samples showed near-zero pathogen abundance, whereas insect samples frequently exceeded 50% and reached 90% relative pathogen abundance. This contrast directly supports the central premise of the sentinel framework; flies enrich and represent pathogen signal(s) environmental sampling alone would miss (H1, supported).

**FIG 1.**
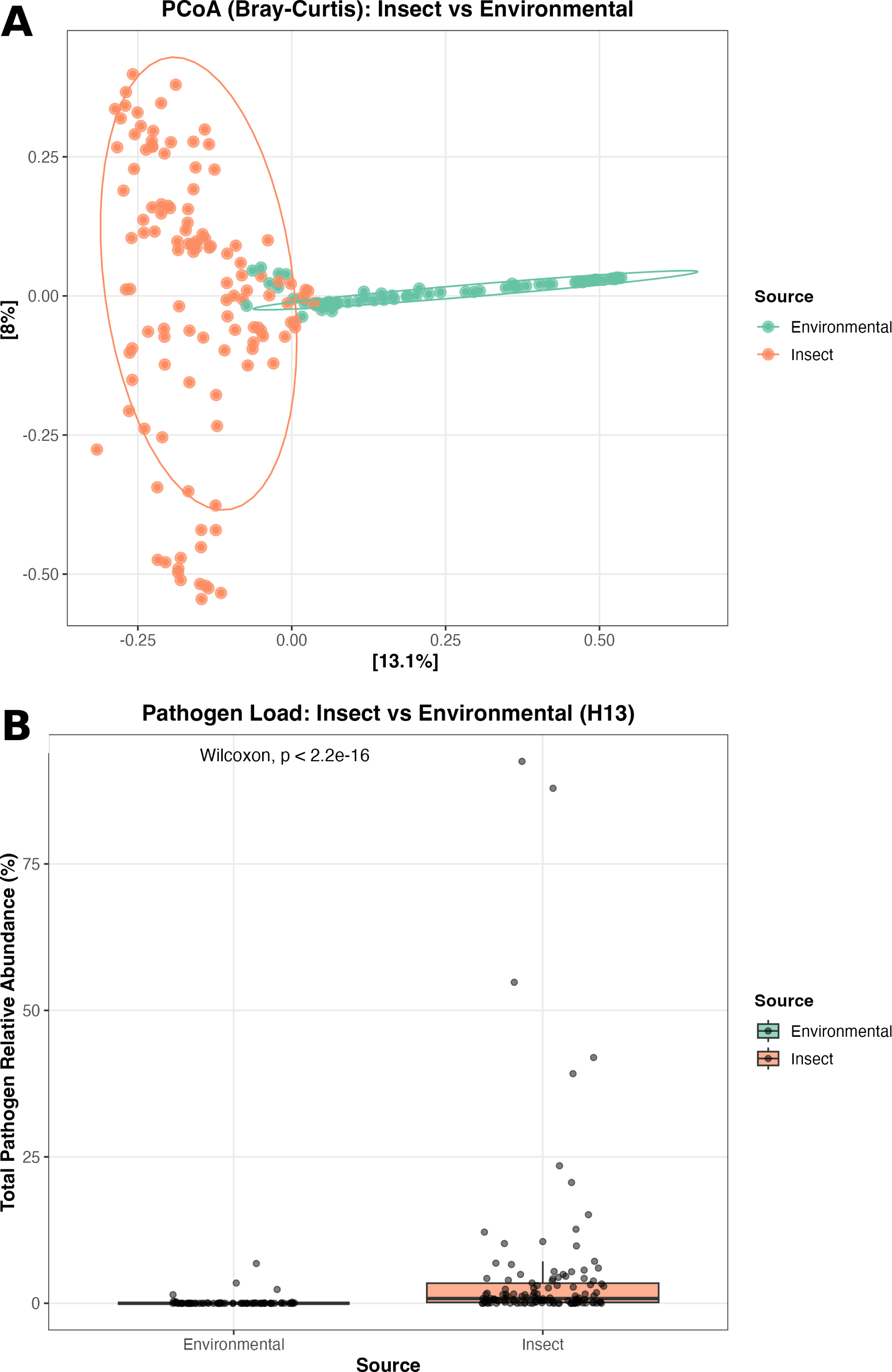
Fly metagenomes form a distinct, pathogen-enriched compartment. (A) Principal-coordinates analysis (PCoA) of Bray–Curtis dissimilarity contrasting insect (orange) and environmental (teal) microbial communities; the first two axes explain 21.1% of variance, and PERMANOVA confirms strong compositional separation (R² = 0.094, F = 19.81, P = 0.001). (B) Total pathogen relative abundance is significantly higher in insect than environmental samples (Wilcoxon P < 2.2 × 10⁻¹⁶); most environmental samples are near zero, whereas insect samples frequently exceed 50%.

### Microbial diversity responds to cattle presence, not to a density gradient

Alpha-diversity patterns dissociated richness from evenness. Observed species richness differed significantly across livestock intensity (Kruskal–Wallis χ² = 19.35, P < 0.001; Hill q0 P<0.01), whereas Shannon (P = 0.516) and Simpson (P = 0.771) diversity did not, indicating that the number of species varied with intensity. At the same time, the evenness of their distribution did not. County-level variation was pronounced across all metrics (Shannon χ² = 23.23; richness χ² = 24.18; Pielou evenness χ² = 27.61; all P < 0.001), reflecting geographic heterogeneity beyond livestock classification. Critically, Jonckheere–Terpstra trend tests detected no monotonic Low→Medium→High trend for any alpha-diversity metric (all P > 0.5), rejecting a simple dose-response model. The pathogen subcommunity behaved similarly; pathogen Shannon diversity (P = 0.574) and richness (P = 0.303) were intensity-invariant. However, total pathogen load differed (P = 0.012) without a monotonic trend (P = 0.996), indicating amplification of specific taxa rather than a graded community shift.

By contrast, the categorical presence of cattle (a county-level classification) was a strong and consistent correlate. Cattle presence was associated with significantly higher Shannon diversity, observed richness, and Pielou evenness (Wilcoxon P < 0.001, 0.042, and < 0.001, respectively). Generalized linear models showed cattle presence predicted Shannon diversity (estimate = 0.769, P < 0.001) and pathogen Shannon diversity, while numeric intensity did not (P = 0.32). Cattle presence and intensity, therefore, act as partially independent observational correlates of community structure rather than as one mediating the other (H3, supported).

### Beta diversity confirms cattle presence as the dominant compositional axis

A combined LivestockIntensity × Cattle model explained 11.6% of Bray–Curtis compositional variation (R² = 0.116, F = 3.84, P = 0.001) and 11.1% of Jaccard variation (P = 0.001), moderate effects typical of microbial-ecology field studies with high environmental heterogeneity. County explained slightly more variation (R² = 0.141, P = 0.001), indicating an independent geographic signal that competes with the livestock classification. Distance-based redundancy analysis confirmed that both intensity (F_2,118_ = 3.24, P = 0.001) and cattle (F = 4.73, P = 0.001) were independently significant after mutual adjustment, with the stronger effect for cattle (Fig. 2), and ANOSIM corroborated this ordering (cattle R = 0.151; intensity R = 0.105). Betadisper was non-significant across intensity levels (F = 0.145, P = 0.874), confirming that the PERMANOVA results reflect genuine location shifts rather than differences in within-group dispersion. For the pathogen subset, the compositional effect was weaker but still significant (R² = 0.057, P = 0.035). The cattle-by-intensity interaction term was not significant (P = 0.54), indicating additive rather than interactive effects.

**FIG 2.**
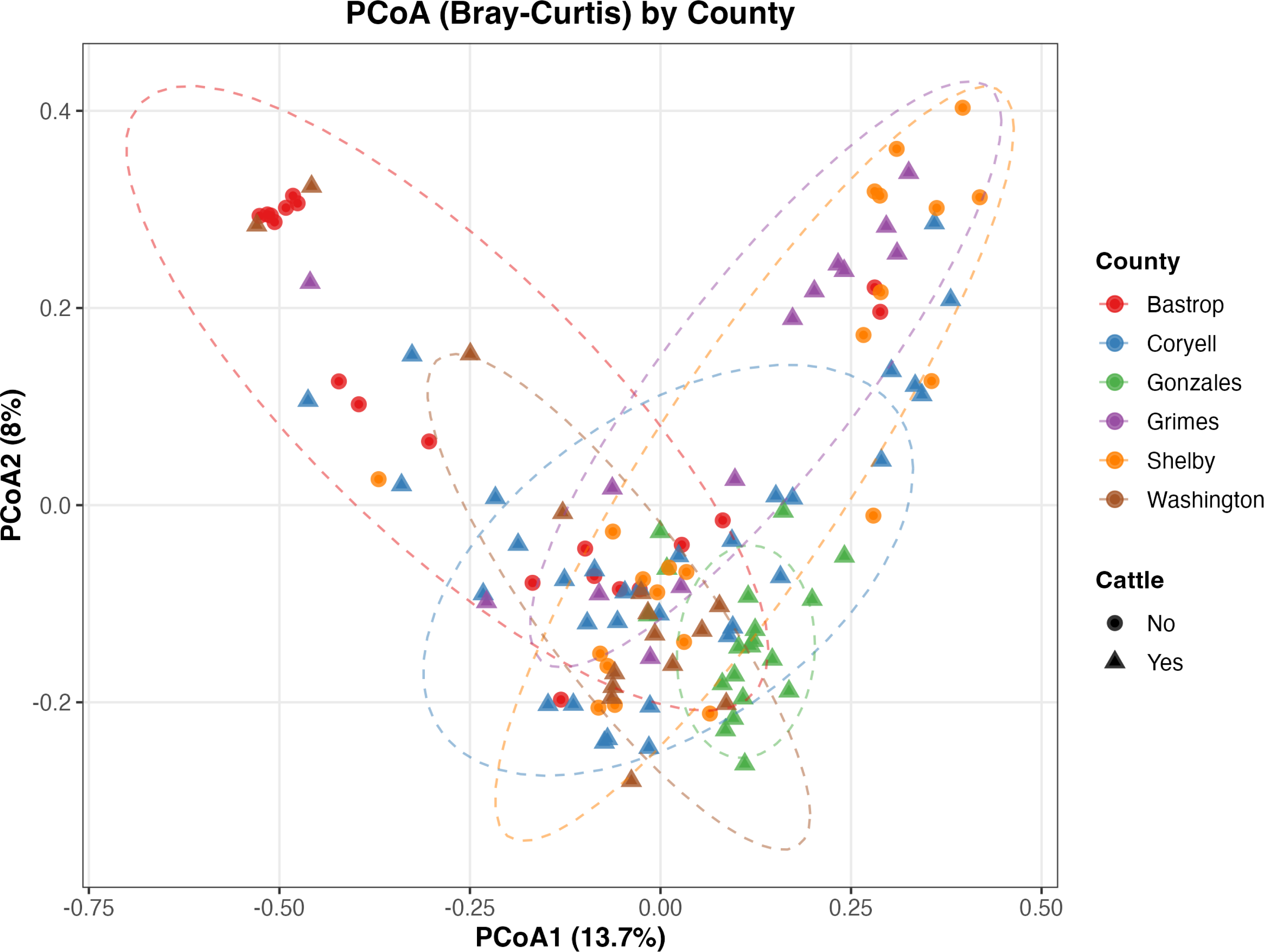
Cattle presence is the dominant driver of community composition. PCoA of Bray–Curtis dissimilarity colored by county and shaped by cattle presence (circles, absent; triangles, present), with 95% confidence ellipses. Distinct clustering of cattle-positive samples (e.g., Washington and Coryell) reflects the categorical cattle signal; db-RDA confirmed cattle (F = 4.73) and intensity (F = 3.24) as independent predictors.

### A gut biomarker anchors the cattle signal

The taxonomic signature of cattle presence was anchored by *Peptostreptococcus russellii*. This strictly anaerobic, hyper-ammonia-producing gut bacterium was the single most robust differentially abundant taxon in the dataset, enriched in cattle-present counties under both ANCOM-BC2 (log-fold change = 2.38, q = 1.27 × 10⁻⁸) and MaAsLin2 (coefficient = 6.44, q = 4.57 × 10⁻⁸; Fig. 3). It was notably the only taxon that passed both the sensitivity-score test and the standard differential-abundance threshold, marking it as statistically robust and highly reproducible. Its detection at high abundance in highly aerobic whole-fly homogenates indicates that flies sample and represent internal core cattle microbiomes in a non-rumen matrix. Conversely, multiple lactic acid bacteria (*Leuconostoc citreum*, *Leuconostoc pseudomesenteroides*, *Enterococcus faecium*) and environmental, plant-associated taxa (*Pantoea agglomerans*, log-fold change = −2.41, q = 2.5 × 10⁻⁵; *Bifidobacterium longum*, log-fold change = −1.94, q = 2.3 × 10⁻⁶) were significantly depleted where cattle were present, consistent with competitive displacement of the background community by ruminant-associated microbiota. MaAsLin2 corroborated these signatures and additionally flagged the arthropod-associated *Rickettsiella grylli* as enriched with cattle presence.

**FIG 3.**
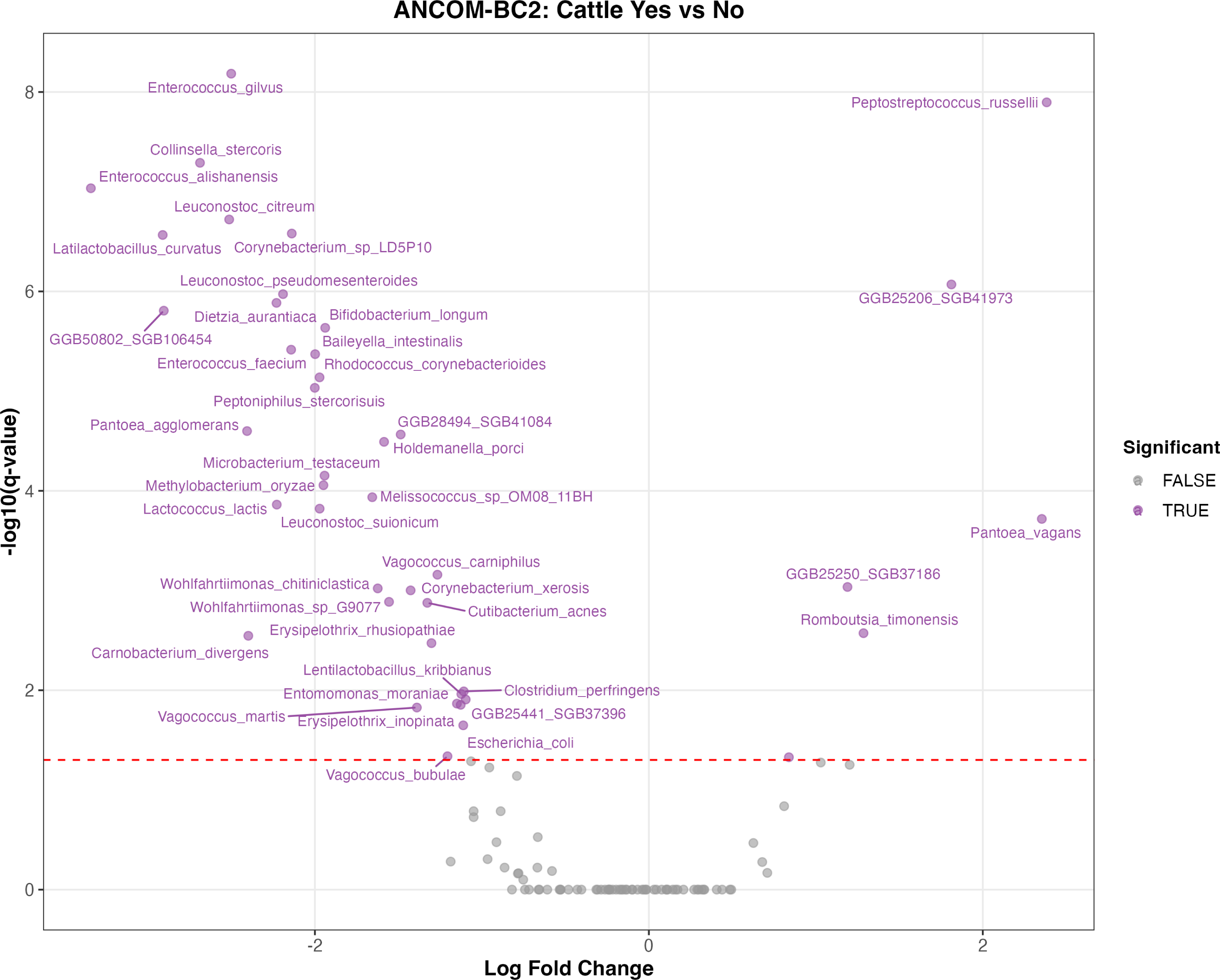
Differentially abundant taxa between cattle-present and cattle-absent environments (ANCOM-BC2 volcano plot). The x-axis shows log-fold change and the y-axis statistical significance (−log10 q). *Peptostreptococcus russellii* is significantly enriched where cattle are present and was the single most robust biomarker in the dataset, whereas many environmental and lactic-acid taxa are enriched where cattle are absent.

### Intensification enriches specific pathogens without community turnover

Although the community showed no smooth dose-response, livestock intensity drove clear taxon-level enrichments. The pathogen *Trueperella pyogenes*, the principal cause of liver abscesses in cattle, showed the strongest intensity effect in the dataset (log-fold change = 2.50, q = 2.62 × 10⁻⁸) and was independently identified as a high-intensity indicator species (Fig. 4). Indicator-species analysis recovered 62 significant taxa across intensity levels: high-intensity environments were marked by livestock-associated organisms (*T. pyogenes*, *Clostridium botulinum*, *Latilactobacillus curvatus*, *Corynebacterium* spp.), whereas low-intensity sites were characterized by insect endosymbionts and fermentation taxa (*Wolbachia pipientis*, *Enterococcus durans*, *Lactococcus petauri*). MaAsLin2 identified 29 significant full-community associations, including enrichment of the arthropod symbiont *Rickettsiella grylli* and *Clostridium botulinum* under livestock pressure.

**FIG 4.**
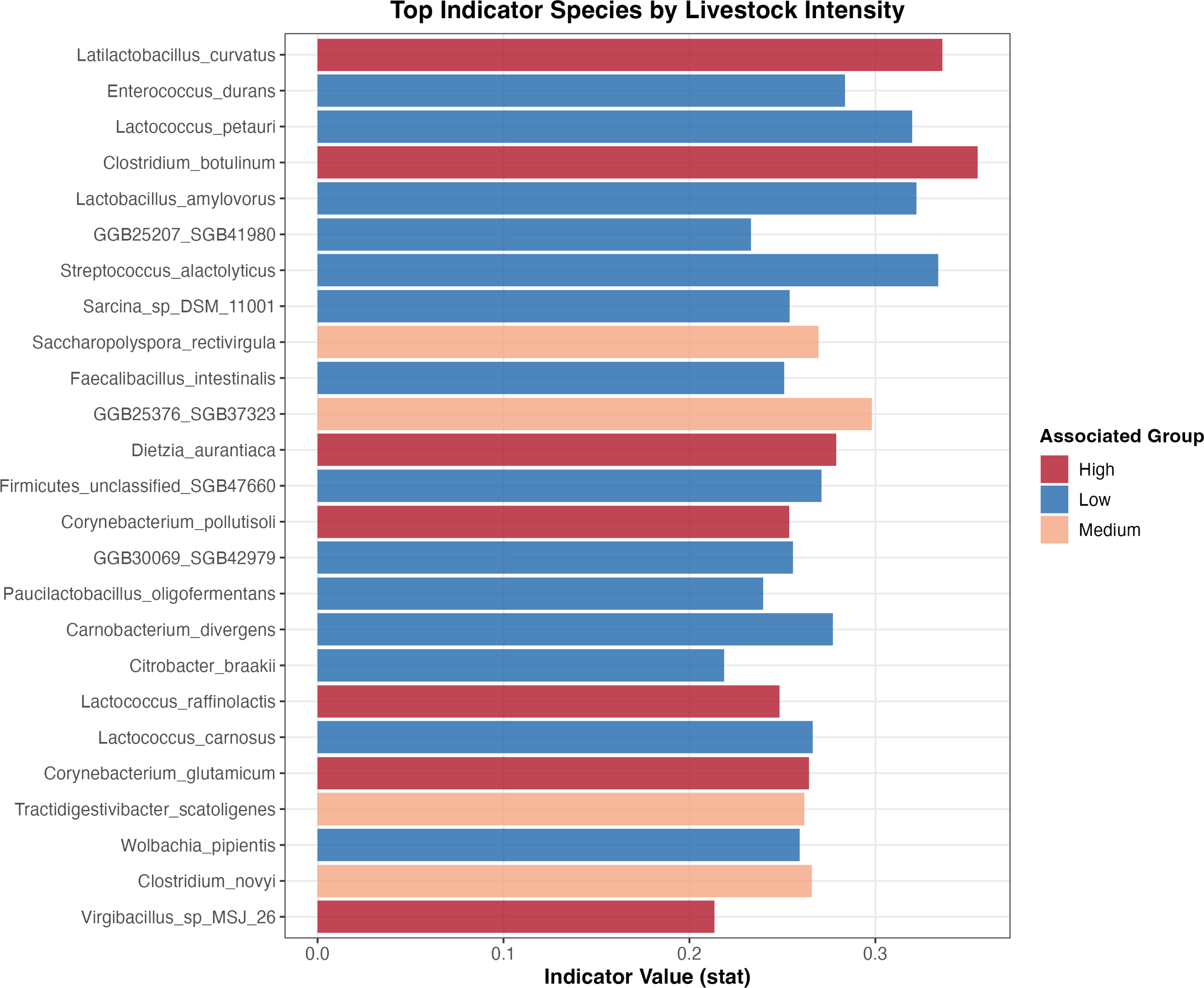
Top indicator species by livestock intensity. Indicator values (stat) for taxa significantly associated with High (red), Medium (tan), and Low (blue) intensity. High-intensity environments are marked by livestock-associated organisms including *Trueperella pyogenes*, *Clostridium botulinum*, and *Latilactobacillus curvatus*; low-intensity sites are characterized by insect endosymbionts and fermentation taxa.

Species richness differed across intensity (P < 0.001), declining at high intensity (Fig. S2), while Shannon, Simpson, and Pielou evenness did not; total pathogen relative abundance also differed (P = 0.012) but non-monotonically, peaking at medium intensity (Fig. S3, S4) rather than rising with livestock pressure. Intensification thus reshapes specific taxa without a community-wide, graded turnover, and conventional diversity indices remain largely insensitive to the change. The cattle pathogen *Chlamydia pecorum*, an obligate intracellular organism of high zoonotic concern, showed a peaked, non-linear response with maximum abundance at intermediate intensity (linear q = 0.005; quadratic q = 0.0013), and *Wolbachia pipientis* showed non-monotonic modulation reflecting shifts in the underlying fly population rather than environmental contact.

### Co-occurrence networks fragment under livestock intensification

Correlation-based co-occurrence networks (Spearman |ρ| > 0.6, P < 0.01) revealed a consistent gradient in network architecture. Low-intensity communities formed the densest, least modular networks (292 edges, density = 0.058, modularity = 0.561); medium-intensity communities were intermediate, and high-intensity communities were the sparsest and most modular (114 edges, density = 0.022, modularity = 0.807). Node-degree distributions shifted accordingly: low-intensity nodes reached degrees as high as 20, whereas high-intensity nodes were dominated by poorly connected taxa (Fig. 5). The progressive loss of density and rise in modularity, characteristic of perturbed ecosystems in which specialist associations replace generalist ones, offers a hypothesis-generating, ecosystem-level descriptor of how livestock pressure reshapes the microbial landscape that flies sample. These correlation networks are descriptive and may be influenced by sample size and prevalence filtering across groups.

**FIG 5.**
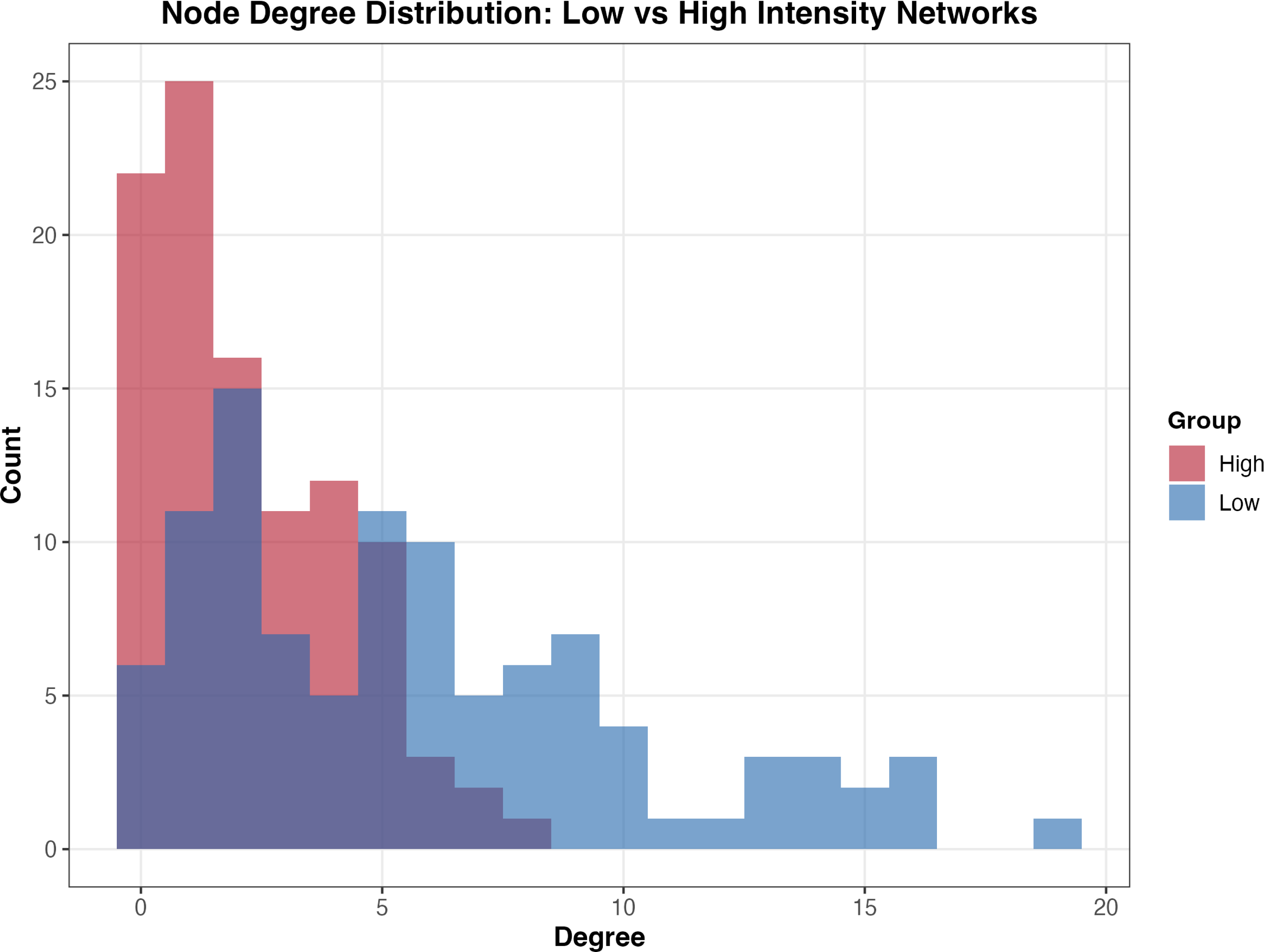
Co-occurrence networks fragment under intensification. Node-degree distributions for Low-(blue) and High-intensity (red) networks. The High-intensity network is skewed toward poorly connected nodes (degree 0–3), while the Low-intensity network reaches degrees as high as 20, indicating a denser, more interconnected community that fragments under livestock pressure (Low density = 0.058, modularity = 0.561; High density = 0.022, modularity = 0.807).

### A narrow pathogen panel fails where whole-community analysis succeeds

A Random Forest model trained on CLR-transformed abundances of a curated 12-pathogen panel classified livestock intensity with only 35.5% out-of-bag accuracy, essentially at parity with a majority-class baseline, and showed heavy bias toward the dominant Medium class (H4, not supported; Fig. S5). Only two pathogen taxa (*Corynebacterium amycolatum*, *Pantoea ananatis*) were significant intensity indicators, and a pathogen-only MaAsLin2 analysis yielded a single significant association. By contrast, whole-community analyses recovered the cattle signal, the *T. pyogenes* intensity signal, dozens of indicator taxa, and the network-fragmentation gradient. The informative content of the fly sentinel therefore lives in the whole microbiome, not in a short list of usual-suspect pathogens.

### Evidence across four focal hypotheses

The four focal hypotheses give a coherent picture (Table 2). Flies form a distinct, pathogen-enriched compartment relative to the co-located environment (H1, supported), and the expectation of a smooth Low–Medium–High dose-response is rejected (H2, not supported). Hence, landscape effects are threshold-like rather than graded. Cattle presence is the strongest landscape correlate across the six counties (H3, supported as an observational association), and a curated 12-pathogen panel cannot substitute for whole-community analysis (H4, not supported), reinforcing rather than weakening the sentinel argument. Together these results indicate that fly metagenomics distinguishes low-impact from livestock-impacted landscapes rather than estimating a quantitative dose-response to animal density.

**TABLE 2.** Evidence summary for the four focal hypotheses.

| ID | Hypothesis | Test / statistic | Decision | Interpretation |
| --- | --- | --- | --- | --- |
| H1 | Fly metagenomes are a distinct, pathogen-enriched compartment relative to co-located environmental samples | PERMANOVA $R^2 = 0.094$,<br>$F = 19.81$, $P = 0.001$<br>(paired within site) | Supported | Cornerstone of the sentinel argument: flies act as a selective filter, not passive mirrors of the environment. |
| H2 | Community structure follows a monotonic Low→Medium→High intensity dose-response | Jonckheere–Terpstra, $P = 0.866$ | Not supported | Landscape effects are threshold-based, not graded: no smooth density gradient. |
| H3 | Cattle presence is associated with community structure (observational, county-level contrast) | GLM Shannon estimate = $0.769$, $P < 0.001$; db-RDA $F = 4.73$, $P = 0.001$; numeric intensity n.s. ($P = 0.32$) | Supported (observational) | Cattle presence is the strongest landscape correlate across the six counties: an association, not a designed contrast. |
| H4 | A curated 12-pathogen panel can substitute for whole-community analysis in classifying landscape context | Random Forest OOB accuracy = $35.5\%$ (≈ majority-class baseline) | Not supported | Narrow panels discard most of the signal; whole-microbiome analysis is required. |

## DISCUSSION

Pooled blow flies (Calliphoridae, 99.6% of specimens) provide a distinct, spatially integrated microbial compartment useful for surveillance of livestock-adjacent landscapes. Our data establish that fly-derived metagenomes are compositionally distinct from co-located environmental matrices and that this compartment encodes reproducible landscape context, particularly cattle presence. They do not establish bacterial viability, transmission directionality, gut colonization versus cuticular carriage, or fine-grained quantification of animal density; claims are accordingly scoped to microbial taxa and pathogen signatures as proxies for livestock-associated exposure contexts.

The compositional separation of fly and environmental microbiomes is the cornerstone of the bioaccumulator argument. If fly microbiomes simply mirrored surrounding soils, waters, and feces, there would be no added value in sampling them. Mechanistically, the fly digestive tract subjects ingested microbes to the peritrophic matrix, reactive oxygen species, and antimicrobial peptides, filtering for taxa biologically compatible with animal hosts and persistent transmission (15). Because many survival-competent organisms are also host-associated pathogens, this selection coincidentally enriches for taxa relevant to zoonotic and AMR risk, so that flies act as signal-enhancing filters rather than noise-adding samplers, isolating the subset of the environmental microbiome with the traits required to colonize, persist, and disseminate.

Cattle presence rather than livestock density emerged as the strongest correlate of community structure, and *Peptostreptococcus russellii* was the most robust biomarker across two independent statistical frameworks. This is exactly the behavior expected of a bioaccumulating sentinel: flies feeding on manure, ocular secretions, and wounds repeatedly contact the rumen-derived bacterial output of cattle, producing a reproducible binary signal that identifies an ecological threshold rather than a graded relationship. The signal is highly sensitive to general livestock contact, reflected in the commensal community, even though amplification of primary zoonotic pathogens may require additional factors. The detection of *Trueperella pyogenes* enrichment near intensive operations is the result with the most direct clinical resonance: liver abscesses are occult lesions typically identified only at slaughter, with U.S. feedlot prevalence of 12–32% and substantial economic cost (16). A non-invasive entomological readout of this burden is precisely the kind of livestock-restricted signal that environmental sampling rarely captures with comparable fidelity.

The absence of a monotonic dose-response should not be read as a failure of the sentinel concept. The most actionable information for a public-health user is the ecological threshold at which a landscape transitions from a low-impact to a livestock-impacted state, and flies reliably distinguish that transition. Species richness in fact declines at high intensity while total pathogen relative abundance peaks at intermediate intensity (Fig. S2, S3), so the shift is taxon-specific rather than a community-wide increase in microbial crowding, and conventional diversity indices remain largely insensitive to it. The progressive fragmentation of co-occurrence networks under intensification adds an ecosystem-structure dimension to this picture, although we offer it as a descriptive, hypothesis-generating result rather than a mechanistic claim about disrupted interactions.

In network ecology, dense, well-connected microbial communities are generally more resilient, whereas a shift toward high modularity with low density is characteristic of perturbed systems in which specialist associations replace generalist interactions (36). The transition we observed, from dense, integrated low-intensity networks to sparse, highly modular high-intensity networks, is consistent with livestock-associated microbes fragmenting the native insect–environment community into more isolated modules. Because these are correlation networks sensitive to threshold choices, prevalence filtering, and group sample sizes, we interpret the topological differences as comparative descriptors rather than mechanistic interaction maps; applying such metrics in the insect–livestock context is uncommon, and we offer the result as a hypothesis-generating observation.

That a curated 12-pathogen panel failed to classify landscape context while whole-community analysis succeeded carries a clear practical implication; future fly-based AMR surveillance should resist reducing monitoring to a small list of usual-suspect pathogens, because doing so discards most of the signal. The corollary is that the value of the fly sentinel scales with analytical breadth. The more of the community that is interrogated, the more landscape context is recovered.

Two further signals illustrate the breadth of information the fly compartment encodes. *Chlamydia pecorum*, an obligate intracellular pathogen of high zoonotic and animal-welfare concern, showed a peaked, non-linear response with maximum abundance at intermediate intensity. Intermediate landscapes may represent a structural “goldilocks zone” in which livestock and wildlife host populations remain connected enough to sustain transmission, whereas highly intensified or sparsely populated environments dampen cycling. That a fastidious organism, routinely missed by culture-based environmental surveillance, was detectable at all in pooled fly metagenomes, and yielded a coherent quantitative pattern, exemplifies the early-warning capability the framework is designed to deliver, with the insect itself acting as the physical livestock–wildlife interface. DNA detection does not establish viability, but it identifies a target for confirmatory sampling.

In parallel, the non-monotonic modulation of the maternally inherited endosymbiont *Wolbachia pipientis* reports not on the environment but on the sentinels themselves. Because *Wolbachia* is host-restricted and vertically transmitted, its abundance in pooled samples reflects the species composition and infection prevalence of the underlying fly population rather than environmental contact. The fly compartment is therefore multi-layered: its exogenous fraction tracks landscape-level pathogen exposure, while its endogenous endosymbiont signal tracks how the vector community itself is restructured by agricultural pressure. Together these layers make filth flies a bioindicator that reports both on the environment and on the sampling organism.

The cumulative weight of evidence supports the bioaccumulator framework. Just as bivalves are used to monitor bioavailable contaminants because they integrate exposure over time, filth flies integrate microbial exposures over the spatial scale of their foraging range, producing a distinct, reproducible compartment detectable by metagenomics. The conceptual contribution of this work is to show, at meaningful sample size, that this compartment is best suited to detecting categorical regime shifts, insect versus environment, cattle present versus absent, low-impact versus livestock-impacted, rather than to estimating continuous animal densities, and that its informative content is maximized when the whole microbiome is analyzed.

Several limitations temper these conclusions. Pooled sampling of 5–10 flies per tube precludes individual-level analyses and obscures sex- and age-related variation. The cross-sectional design captures a single seasonal window. Family-level morphological identification may mask species differences in vector competence. Livestock intensity and cattle presence were assigned at the county scale (two counties per intensity tier), so these landscape factors are confounded with county-level differences in land use, climate, and vegetation and are interpreted as observational associations across six counties rather than designed contrasts. Sequencing depth limits detection of rare pathogens and metagenomic DNA cannot distinguish viable from relic organisms. The 1–2 km foraging radius is a contextual rationale from prior mark–recapture work (7), not a parameter estimated here, and roadside sampling is best understood as a pragmatic sentinel-deployment setting rather than a substitute for on-farm sampling.

The most natural extensions are distance-decay transect designs to quantify the spatial scale over which the sentinel signal persists, longitudinal sampling to resolve seasonal dynamics, paired culture-based validation to establish viability for key signals such as *T. pyogenes* and *C. pecorum*, and targeted characterization of the ARG mobilome within fly gut metagenomes to assess horizontal-transfer potential. Extending the framework to additional vector taxa would broaden it into a multi-vector view of the agricultural resistome. Compared with point-source environmental sampling, entomological metagenomics offers higher information density per sample, lower per-coordinate cost, and a One Health relevance that soil and water cannot match: flies occupy the interfaces among livestock operations, surrounding environments, and human spaces, making them defensible integrators for landscape-scale microbial surveillance.

## MATERIALS AND METHODS

### Study design and site selection

Using county-level livestock-distribution data from USDA-NASS (2019), six Texas counties were stratified into three livestock-intensity tiers: Low (Grimes, Washington), Medium (Bastrop, Coryell), and High (Gonzales, Shelby). Livestock intensity was assessed as the total number of livestock reported in a county, whereas livestock diversity considered species (poultry, cattle, swine etc). The chosen counties were as closely matched in area as possible (Range: 1608 km^2^ – 2766 km^2^, with CV of 19%, compared to statewide average of 64%). The livestock counts distribution by county was divided into to the top, middle, and lower third, and sites within 320 km (by car) from College Station, Texas, were selected for inclusion in the study. Cattle presence or absence was applied based on visual confirmation of cattle within a 5km radius of each sampling site. Within each county, a randomly selected epicenter anchored four surrounding sampling sites positioned ∼15–16 km away, restricted to publicly accessible roadways for safety. Each county was sampled in two collection events. At each site, three two-meter steel U-posts were installed ∼33 m apart along a linear transect and georeferenced. Co-located environmental samples, standing water or runoff, surface soil and decomposing plant matter, and wildlife or livestock feces, were collected near each post to characterize the local background resistome.

### Insect collection and laboratory processing

Modified RESCUE! reusable traps were baited with sweetened, fermented rotted chicken liver in vented cups that emitted odor while preventing contact between flies and bait (17). Traps were operated from 10:00 to 17:00 on days with temperature ≥15.5 °C, minimal rain forecast, and wind <4.47 m/s, upon removal traps were placed on ice for transport. Captured Diptera were chilled to 4 °C, and target taxa (*Musca domestica*, Calliphoridae spp.) were identified morphologically and removed; non-target insects were recorded but excluded. Sub-samples of 5–10 flies matched by Family or species and trap location were pooled and homogenized (BioSpec Mini-Beadbeater-96). Total genomic DNA was extracted with the QIAamp DNA Microbiome Kit (host-depletion optimized) for insects and QIAGEN PowerFecal Pro Kits for environmental matrices. Libraries were sequenced at the Texas A&M AgriLife Genomics and Bioinformatics Service, targeting 5–7 Gbp per sample.

### Bioinformatics

Reads were quality- and adapter-filtered, assembled de novo with MEGAHIT (18), and binned with MetaBinner and MetaBAT 2 to recover metagenome-assembled genomes (19); genomes were annotated with Prokka (20). Assembled contigs were screened for antimicrobial-resistance genes and virulence factors using ABRICATE (21) against the ResFinder (22) and VFDB (23) databases. Species-level taxonomic profiles were generated from quality-filtered reads with MetaPhlAn 4 (24), and functional content was mapped to KEGG (25).

### Statistical analysis

Analyses were performed in R v4.4 using phyloseq for data management (26). Environmental samples were excluded from primary insect analyses (122 retained); taxa present in <5% of samples or with mean relative abundance <0.01% were removed, yielding 254 taxa. For compositional methods requiring integer input, relative abundances were converted to pseudo-counts (proportions × 10⁶, rounded), and downstream differential-abundance estimates are accordingly framed in relative rather than absolute terms. A curated list of 21 clinically and veterinary-relevant pathogen species spanning the genera *Enterococcus*, *Staphylococcus*, *Providencia*, *Chlamydia*, *Bartonella*, *Brucella*, *Pantoea*, *Clostridium*, and *Corynebacterium* was compiled and annotated for Gram stain, zoonotic potential, AMR concern, and primary habitat; 12 of these were detected and retained for pathogen-specific analyses, and the remaining 242 taxa were treated as non-pathogen sentinels.

Alpha diversity (observed richness, Shannon, Simpson, inverse Simpson, Pielou evenness, Hill numbers q = 0, 1, 2) was computed in vegan (27) and compared with Kruskal–Wallis (intensity), Wilcoxon (cattle), and Jonckheere–Terpstra trend tests; GLMs tested independent effects of intensity and cattle presence. Beta diversity used Bray–Curtis and Jaccard distances with PCoA and NMDS ordination, PERMANOVA (adonis2, 999 permutations), ANOSIM, betadisper, db-RDA, and variance partitioning. Differential abundance was assessed with ANCOM-BC2 (28) and MaAsLin2 (29), retaining taxa at q < 0.05. Random Forest classification and regression used CLR-transformed abundances (ranger; 1,000 trees, permutation importance) (30). Indicator-species analysis used multipatt (point-biserial coefficient, 999 permutations) (31). Co-occurrence networks (top 100 taxa plus all pathogens; Spearman |ρ| ≥ 0.6, P < 0.01) were built and characterized for density, transitivity, average degree, and Louvain modularity in igraph (32), overall and stratified by intensity. Mediation and structural-equation models (mediation, lavaan) were used as descriptive decompositions under stated assumptions, not as causal claims, given the observational design (33, 34). Insect-versus-environment contrasts used all 195 samples with Wilcoxon tests and Bray–Curtis PERMANOVA. The complete pipeline is available as a unified R script.

## ACKNOWLEDGMENTS

We thank the Kaufman laboratory (Texas A&M Entomology) for assistance with sampling design, identification, and processing; the Taylor laboratory (Texas A&M Animal Science); and current and past members of the Athrey laboratory (Texas A&M Poultry Science) for sampling, laboratory processing, and support. Sequencing was performed by the Texas A&M AgriLife Genomics and Bioinformatics Service. The work was funded by the Vector Seed Grant Program from Texas A&M AgriLife Research to GA, PEK, and TMT. Alicia Montemayor was also supported on a Graduate Teaching Assistantship from the Department of Poultry Science. The authors declare no competing interests.

## Data Availability Statement

Raw shotgun metagenomic sequence reads generated in this study will be deposited in the National Center for Biotechnology Information (NCBI) Sequence Read Archive (SRA) and will be publicly released upon acceptance. During peer review, the sequence reads and associated sample metadata are available from the corresponding author upon request. Curated species-level abundance tables and the complete analysis pipeline (a unified R script) that support the findings of this study are available from the corresponding author upon request.

## AUTHOR CONTRIBUTIONS

G.A. T.M.T., and P.E.K. conceived and designed the study. P.E.K. advised on trap design, fly collection, and morphological identification, and T.M.T. contributed to study design and provided resources. A.M. performed field sampling, laboratory processing, DNA extraction, and the bioinformatic and statistical analyses. G.A. supervised the project and acquired funding. A.M. and G.A. wrote the manuscript, and all authors reviewed, edited, and approved the final version.

## ETHICS

This study did not involve human subjects, human data or tissue, or experiments on live vertebrate animals. Sampling was limited to synanthropic insects and environmental matrices (soil, water, plant material, and naturally deposited feces) collected from publicly accessible roadsides; accordingly, Institutional Review Board or Institutional Animal Care and Use Committee approval was not required.

